# The compound of baicalein, wogonin, and oroxylin-A inhibits EMT in A549 cell line via PI3K/AKT-TWIST1- glycolysis pathway

**DOI:** 10.1101/2021.05.13.443958

**Authors:** Hui-Juan Cao, Wei Zhou, Xiao-Le Xian, Shu-Jun Sun, Pei-Jie Ding, Chun-Yu Tian, Fu-Ling Tian, Chun-Hua Jiang, Ting-Ting Fu, Shu Zhao, Jian-Ye Dai

## Abstract

Non-small cell lung cancer (NSCLC) is a worldwide disease with high morbidity and mortality, which is most derived from its metastasis. Some researches show that the epithelial-mesenchymal transition (EMT) process promotes lung cancer cells migration and invasion, leading to NSCLC metastasis. Total Flavonoid Aglycones Extract (TFAE) isolated from *Scutellaria baicalensis* was reported to inhibit tumor growth and induce apoptosis. In this study, we found that baicalein, wogonin, and oroxylin-A were the active compounds of TFAE. After reconstructing with these three compounds (baicalein (65.8%), wogonin (21.2%), and oroxylin-A (13.0%)), the reconstructed TFAE (reTFAE) present inhibitory effect on the EMT process of A549 cells. Then, bioinformatic technology was employed to elucidate the potential pharmacodynamic mechanism network of reTFAE. We identified the relationship between reTFAE and PI3K/Akt signaling pathways, with TWIST1 as the key protein. LY294002, the inhibitor of PI3K/Akt signaling pathway, and knock-down TWIST1 could significantly enhance the efficacy of reTFAE, with increasing expression of epithelial markers and decreasing expression of mesenchymal markers in A549 cells at the same time. Furthermore, stable isotope dimethyl-labeled proteomics technology was conducted to complement the follow-up mechanism that the EMT-inhibition process may be realized through glycolysis pathway. In conclusion, we claim that TWIST1-targeted flavonoids could provide a new strategy to inhibit EMT progress for the treatment of NSCLC.

## 1. Introduction

Lung cancer is a worldwide disease with high morbidity and mortality. According to World Health Organization[1], there were 2.207 million new lung cancer patients worldwide in 2020, only less than breast cancer; and lung cancer ranked first in the global causes of death. Among the histological subtypes of lung cancer collectively, non-small cell lung cancer (NSCLC) accounts for 80%-85%; and most lung cancer patients are diagnosed at advanced stages, so traditional chemotherapy and radiotherapy have limited efficacy[2]. NSCLC microenvironment has limited nutrition, so cancer cells often obtain the energy and substances required for proliferation and cell growth through the “Warburg Effect” glucose metabolism changes[3, 4], that is, cancer cells tend to produce energy through glycolysis rather than oxidative phosphorylation[5-7]. However, the metabolic change leads to further tumor cell growth, aggravation of epithelial-mesenchymal transition (EMT) process and resistance to treatment. During the treatment of NSCLC, 40% of patients have metastases[8], and the main biological process is EMT[9]. In EMT process, cells gradually become spindle-shaped, losing the unique epithelial characteristics, intercellular adhesion and cytoskeletal structure of epithelial tissue, and possessing polarity, individual migration ability and invasion ability[9]. Cellular EMT process is related to TGFR, Wnt/β-Catanin, PI3K/Akt, Ras, MAPK, NF-κB pathways, and the transcription factors involved are TWIST (TWIST1 and TWIST2), Slug, ZEB (ZEB1 and ZEB2), Snail et al[10, 11].EMT transcription factors promote the development of drug resistance[12, 13]. In the process of EMT, the activated PI3K/Akt pathway promotes glucose uptake and glycolysis[14, 15], and phosphorylated Akt can phosphorylate TWIST1, then up-regulate the TGFβ-Smad pathway, which further promotes the EMT process [16]. TWIST1 was known to bind to the E-box sequence of E-cadherin and inhibited the expression of E-cadherin to induce EMT in a variety of tumors[17-19].Inhibition of TWIST1 was known to induce growth inhibition and apoptosis of EGFR-mutant NSCLC cells[20].

Some herbals and their extracts have been reported to have therapeutic effects on cancer, including *Scutellaria baicalensis* [21, 22], whose main components are flavonoids. Total Flavonoid Aglycones Extract (TFAE) isolated from *Scutellaria baicalensis* was reported to inhibit tumor growth and induce apoptosis, in which the main components, baicalein, wogonin, and oroxylin-A accounts for 41.4%, 13.3%, and 8.2%, respectively [23]. This inspired us to reconstruct a compound combination with baicalein, wogonin, and oroxylin-A.

In this study, we reconstructed TFAE with baicalein (65.8%), wogonin (21.2%), and oroxylin-A (13.0%) and named it as reTFAE (Fig.1A), and found that the reTFAE could significantly inhibit EMT of A549 cells. We constructed the TFAT-PI3K/AKT-TWIST1 correlation in inhibiting EMT progress, and revealed that this phenomenon may be related to glycolysis pathway through stable isotope dimethyl-labeled proteomics technology. Therefore, we prove that TWIST1 may be a potential target to inhibit the EMT process of lung cancer, and hope to explore a new combined medication strategy through the compatibility study of active compounds in herbal extracts.

**Fig.1.**
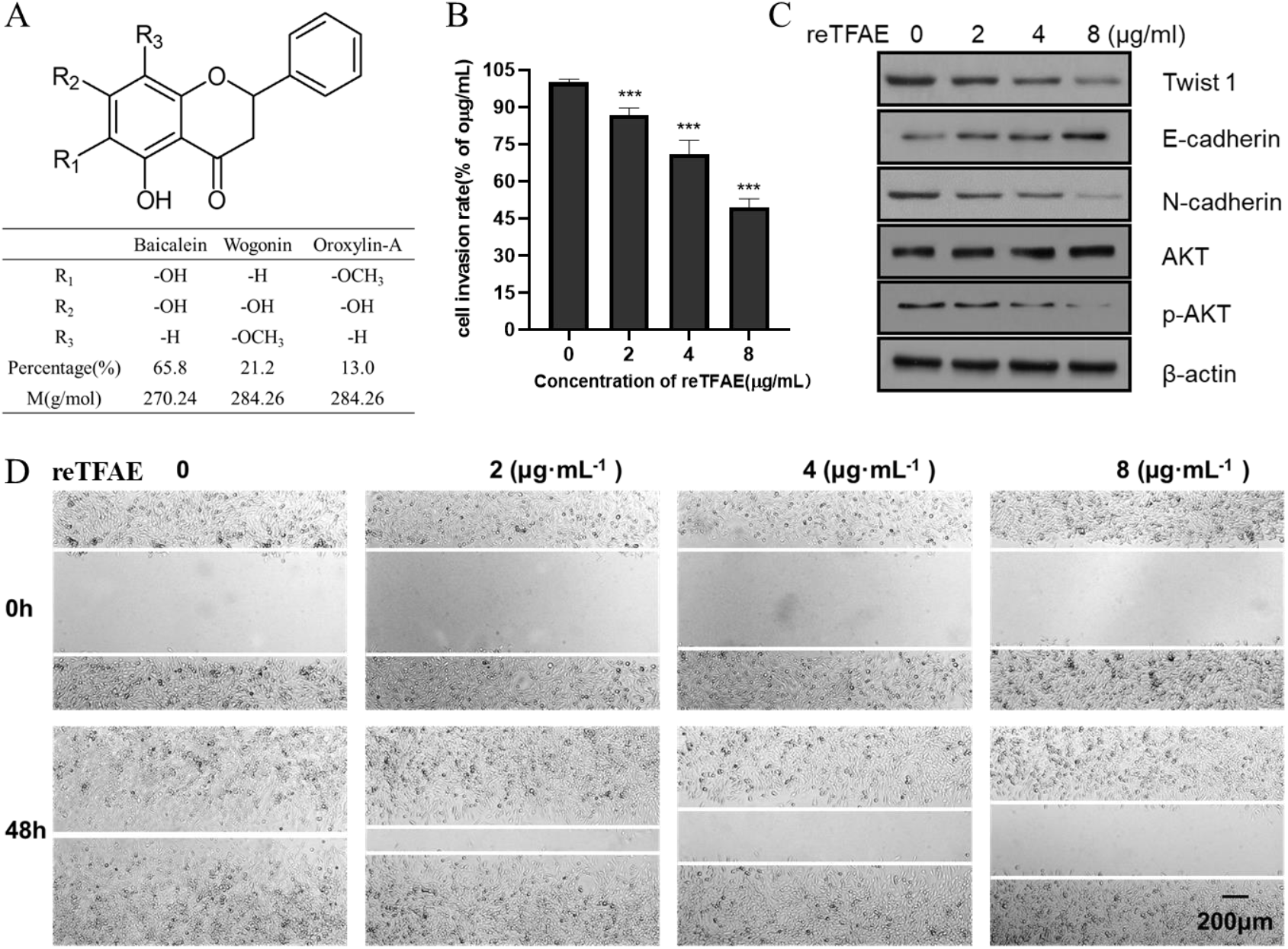
reTFAE inhibited EMT process of A549 cells. (A) The structure and proportion of the main components of reTFAE. (B) The invasion-inhibiting ability of reTFAE on A549 cells. (C) Effects of different concentrations of reTFAE on the expression of EMT-related proteins. (D)reTFAE inhibits A549 cell migration. Statistical differences were determined by a two-sided Student’s t-test. Compared with 0µg/mL reTFAE, \*\*\**P* < 0.001; \*\**P* < 0.01; \**P* < 0.05.

## 2. Materials and Methods

### 2.1. Cell culture

The A549 cells were obtained from the National Collection of Authenticated Cell Culture, and were cultured at 37 °C under 5 % CO_2_ in Dulbecco’s modified Eagle’s medium (DMEM; Sigma, D6429) supplemented with 10% (vol./vol.) fetal bovine serum (FBS; Gibco, 10270-106), and 1 % (vol./vol.) penicillin-streptomycin (PS; Gibco, 15140-122). Culture medium was refreshed and the cells were passaged when the confluency was 80%.

### 2.2. Preparation of cisplatin, LY294002 and reTFAE

4mg/mL cisplatin stock solution, 20mM LY294002 stock solution and 128mg/mL reTFAE solution were prepared by DMSO and were used for the later experiments. For reTFAE, the three main components and their mass percentages are baicalein (65.8%, Sichuan Weikeqi Biological, wkq-00289), wogonin (21.2%, Sichuan Weikeqi Biological, wkq-00245), and oroxylin-A (13.0%, Sichuan Weikeqi Biological, wkq-00180). We weighed the amount of each component according to their mass percentage, then mixed the three compounds, and dissolved them to the desired volume with DMSO.

### 2.3. Wound healing assay and cell invasion assay

For wound healing assay: 5×10^6^ A549 cells were plated in six-well plates. 7 hours later, the cells formed a single layer with nearly 100% confluency. Confluent cell layers were scratched using 200μL tips to generate wounds with approximately 800 µm width. Cells were washed with PBS for 3 times to remove detached cells and debris. Cells were incubated with complete medium with or without reTFAE, Cisplatin, or LY294002 for 24 h. At 0 and 24h after scratching, 3 fields of view for each well were select randomly to take pictures. Image J software were used to scan the scratch areas and to calculate the wound healing area. Wound healing rate= (wound area at 0h-wound area at 24h)/ wound area at 0h *100%.

For cell invasion assay: the transwell containing polycarbonate membranes with 8µm pore size was precoated with 50µL of Matrigel(1.0mg/mL) and incubated at 37°C overnight. 1.0*10^6^ A549 cells and reTFAE (2, 4, 8 µg/mL) in serum-free DMEM medium were plated in the upper chamber, followed by 150µL of serum-supplemented DMEM medium in the lower chamber. After 24h of incubation, the upper chamber was taken out and washed by wash buffer for 2 times, then was transferred to test board containing 100µL Cell Dissociation Solution/Calcein-AM. After 1h of incubation, the fluorescence value was detected at 485nm excitation wavelength and 520nm emission wavelength. Cell invasion rate= (number of invaded cells in experimental wells/ number of invaded cells in control wells) *100%.

### 2.4. Western blot

The A549 cells under different conditions were collected after washed by PBS for three times and centrifugation at 3000 rpm for 3 minutes. The cells were lysed in RIPA Lysis Buffer (CWBIO, CW2333S) containing EDTA-free cOmplete(Roche, 4693132001) with sonication on ice. The cell lysates were collected by centrifugation (150*100rpm, 20min) at 4°C. The protein concentration was detected by Pierce™ BCA Protein Assay Kit (Thermo Fisher Scientific, 23225) and adjusted to 2mg/mL.20µg of the lysates were separated by 10% SDS-PAGE and then transferred onto a PVDF membrane in an ice bath at 300mA for 1h. The membrane was blotted with 5% nonfat milk in TBST at RT for 1h and then incubated with the following antibodies at 4°C overnight: TWIST1(Cell Signaling Technology, 46702S), E-cadherin (Cell Signaling Technology, 3195), Vimentin (Cell Signaling Technology,5741), N-cadherin (Santa, Sc-59987), AKT (Cell Signaling Technology, 4685), p-AKT (Cell Signaling Technology, 4060), β-Actin (Proteintech, 20536-1-AP). GAPDH (Proteintech, 10494-1-AP). Then the membrane was incubated with secondary antibodies at RT for 1h. The secondary antibodies are as follows: Goat anti-Rabbit IgG (Proteintech, SA00001-2), Goat anti-mouse IgG (Proteintech, SA00001-1). Finally, the immunoreactive bands were visualized by the Clarity™ Western ECL Substrate (BIO-RAD,1705061) using the Tanon4600 Automatic chemiluminescence image analysis system (Tanon, Shanghai, China).

### 2.5. Transfection

6.0*10^5^ A549 cells were plated in six-well plates. 7 hours later, the cells with nearly 70% confluency were transfected with TWIST1 siRNA. The siRNA sequences were as follows: TWIST1(sense:5’-CAAGAUUCAGACCCUCAAGTT-3’, antisense:5’-CUUGAGGGUCUGAAUCUUGTT-3’); GAPDH (sense:5’-AATGGGCAGCCGTTAGGAAA -3’, antisense: 5’-TGAAGGGGTCATTGATGGCA -3’). A siRNA nonspecific control was also used in this study. SiRNA sequences were designed and synthesized by GenePharma (Shanghai, China). The interference was performed based on the protocol of Lipofectamine^®^ 3000 (Thermo Fisher Scientific). The knockdown efficiency was examined by qRT-PCR.

### 2.6. Quantitative real-time PCR (qRT-PCR)

Total RNA was extracted and purified with Eastep® Super Total RNA Extraction Kit (Promega, LS1040) and used as the cDNA synthesis template for reverse transcription with the RevertAid First Strand cDNA Synthesis Kit (Thermo Scientific, k1622). The reverse transcription conditions were as follows: 42°C 30min; 85°C 10min. qRT-PCR was performed by MX3000P(Agilent) using UltraSYBR Mixture (Low ROX) (CWBIO, CW2601), reaction conditions were :95°C 3min; 95°C 15s; 55°C 30s, 72°C 30s; 40 cycles. The primers were as follows: TWIST1(forward: 5’-TCGGACAAGCTGAGAGCAAGATTCA-3’, reverse: 5’-TCCATCCTCCAGACCGAGAAGG - 3’), GAPDH (forward: 5’- CATGAGAAGTATGACAACAGCCT-3’, reverse: 5’-AGTCCTTCCACGATACCAAAGT-3’). CT values were used to evaluate the relative mRNA expression by 2^-ΔΔCt^ method and β-Actin served as an internal control. The primers were designed and synthesized by GenePharma (Shanghai, China).

### 2.8. Protein in-solution digestion and dimethyl labeling

A549 cells were grown to 80% confluence in 15cm dish. A549 cells were treated with 64µg/mL reTFAE for 24h. The cells were collected after washed by PBS for three times and centrifugation at 3000 rpm for 3 minutes. The cells were lysed in 0.1% Triton X-100(Sigma-Aldrich, T8787)-100mM TEAB (Sigma-Aldrich, T7408) containing EDTA-free cOmplete with sonication on ice. The cell lysates were collected and the protein concentration was detected by Pierce™ BCA Protein Assay Kit. 10µL cell lysates(3mg/mL) were reacted with 30µL 8M urea and 2µL 200mM DTT at 65°C for 15min in the dark. Then 2µL 400mM iodoacetamide was added to react in light for 30min at 35°C. 2µL 200mM DTT was added to react with the remaining iodoacetamide in light for 15min at 65°C. 100µL 100mM TEAB, 2µL 0.2µg/µL trypsin (Promega, V528A) and 1.5µL 100mM CaCl_2_ were added and the trypsin digestion was performed at 37°C overnight. For dimethyl labeling: for the control(“light”) group, the peptides were reacted with 6µL 4% CH_2_O (Sigma-Aldrich, F1635) and 6µL 0.6M NaBH_3_CN (Sigma-Aldrich, 42077); for the reTFAE (“heavy”) group, the peptides were reacted with 6µL 4% ^13^CD_2_O (Sigma-Aldrich, 596388) and 6µL 0.6M NaBD_3_CN(Sigma-Aldrich,190020). The reaction was incubated at 22°C for 1h. The reaction was quenched by 24µL 1% ammonia and 12uL formic acid. The “light” and “heavy” samples were combined and the desalination was performed by Pierce™ C18 Tips (Thermo Scientific,87782).

### 2.8. LC-MS/MS analysis

Samples were analyzed by LC-MS/MS on EasyNano-LC and Q Exactive series Orbitrap mass spectrometers (Thermo Fisher Scientific). In positive-ion mode, full-scan mass spectra were acquired over the m/z ratio from 350 to 1800 using the Orbitrap mass analyzer with mass resolution of 7000. MS/MS fragmentation is performed in a data-dependent mode, and the TOP 20 most intense ions are selected for MS/MS analysis with a resolution of 17500 under HCD’s collision mode. The isolation window was 2.0 m/z units, the default charge was 2+, the normalized collision energy was 28%, the maximum IT was 50ms, and dynamic exclusion, was 20.0s. Under +57.0215 Da’ s cysteine modification, the LC-MS/MS data was analyzed by ProLuCID. The isotopic modifications were set as static modifications on the N-terminal of a peptide and lysins, and 28.0313 and 34.0631 Da were for light and heavy labeling, respectively. The CIMAGE software was applied for quantitation, proteins with an average ratio(light/heavy) above 1.5 were selected for further KEGG analysis.

### 2.9. Network pharmacology analysis

In this study, the common mechanism of reTFAE in the treatment of lung cancer was studied based on network pharmacology. The active ingredients and related targets of reTFAE were integrated from TCMSP, BATMAN-TAM, STP and Pubchem databases. The standard names of these targets were united by UniProt database. Targets of lung cancer were enriched through GeneCards, NCBI(Gene), Therapeutic Target Database, and DisGeNET(v7.0) databases. Then the intersection targets of reTFAE and disease were obtained. The STRING network and the Cytoscape 3.7.2 were used to construct Protein-Protein-Interaction (PPI) network, the DAVID database was used to perform KEGG analysis. Then Cytoscape 3.6.1 was used to build “Ingredient-Target-Signal Pathway” network.

### 2.10. Statistical analysis

Student’s t-test was used to compare experimental data. We analyzed the data in GraphPad Prism (GraphPad Software), using the unpaired, two-tailed t-test module. Statistical significance was considered when a p value was below 0.05. \**p*< 0.05; \*\**p* < 0.01; \*\*\**p* < 0.001. N.S. not significant.

## 3. Results

### 3.1. reTFAE inhibited EMT processes of A549 cells

To explore reTFAE’s effect in the EMT process of lung cancer, A549 cells were treated with reTFAE (2, 4, 8 µg/mL) for 24 hours. Compared to 0µg/mL reTFAE, the invasion rate of cells treated by reTFAE concentration-dependently decreased (Fig.1B). In addition, reTFAE dose-dependently inhibited the expression of TWIST1, p-AKT and N-cadherin proteins, while increased the expression of E-cadherin (Fig.1C). Compared with the wound area at 0 hours, the wound were dose-dependently healed by reTFAE(Fig.1D), which means reTFAE inhibited the migration of A549 cells.

### 3.2. reTFAE inhibited EMT process via PI3K/Akt pathway

The above results indicated that reTFAE exerted a concentration-dependent inhibitory effect on the EMT process of A549 cells. To explore the connection between reTFAE and lung cancer, network pharmacology analysis was performed. Three active ingredients and 65 related targets of reTFAE were integrated from TCMSP, BATMAN-TAM, STP and Pubchem databases. The targets of lung cancer were enriched through GeneCards, NCBI(Gene), Therapeutic Target Database, and DisGeNET(v7.0) databases. Venn diagram showed the intersection of 62 reTFAE and lung cancer targets (Fig.2A). STRING network and Cytoscape 3.7.2 were used to construct Protein-Protein-Interaction (PPI) network (Fig.2B), AKT1 was the central protein. DAVID database was used to perform KEGG analysis, reTFAE regulated pathways with P<0.01 were as follows: TNF signaling pathway, p53 signaling pathway, PI3K/Akt signaling pathway, VEGF signaling pathway, HIF-1 signaling pathway, MAPK signaling pathway (Fig.2C). After literature researching, PI3K/Akt signaling pathway attracted our attentions, which could significantly activate tumor metastasis and directly regulate the expression of TWIST1, one of the key markers in EMT [16]. To investigate whether reTFAE affects the EMT process of A549 cells by acting on PI3K/Akt signaling pathway, we incubated A549 cells with the PI3K/Akt pathway inhibitor, LY294002. In this part, cisplatin was as positive control.

**Fig.2.**
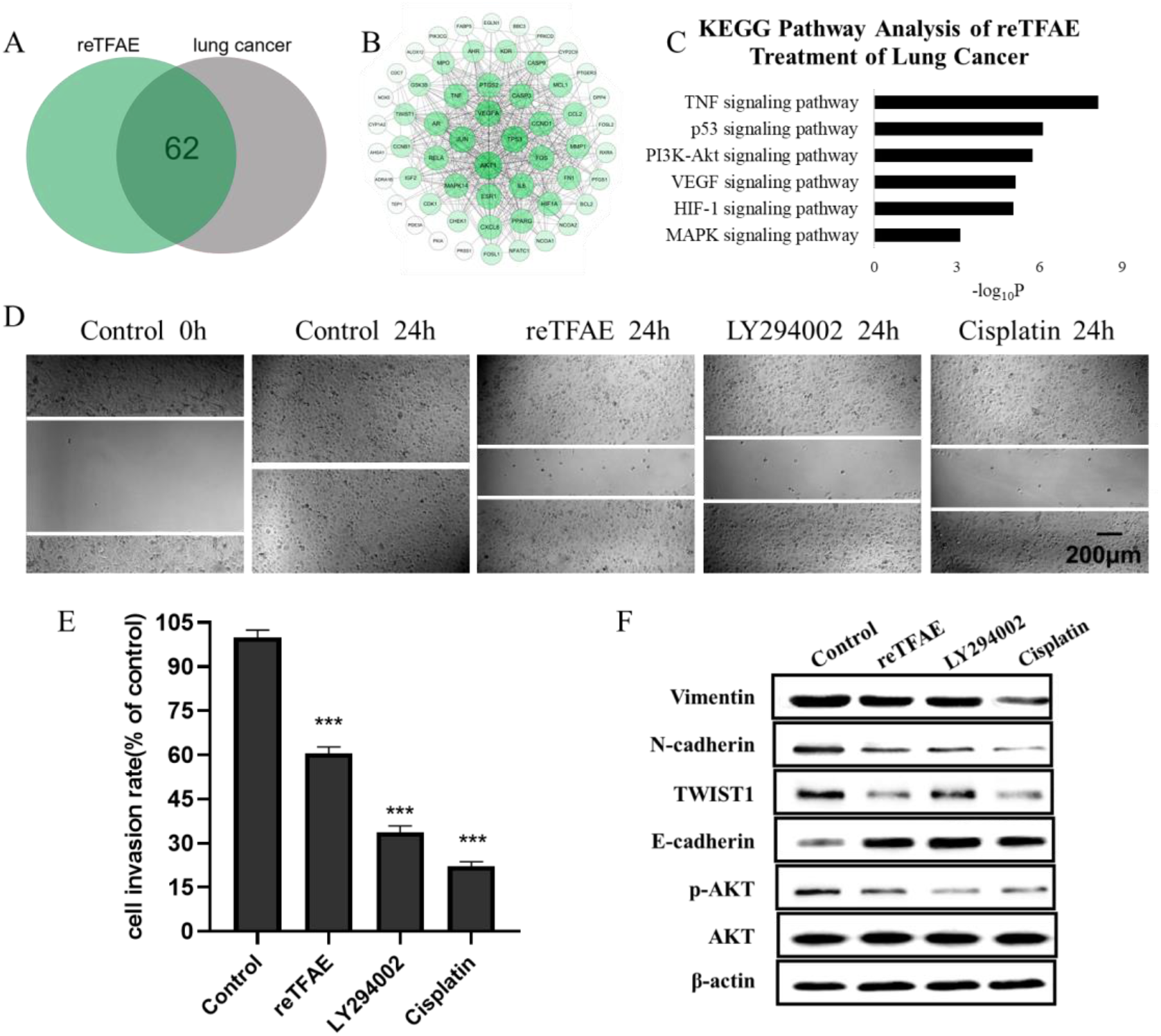
reTFAE inhibited PI3K/Akt pathway to ameliorate EMT process of A549 cells. (A) Intersection protein targets of reTFAE and lung cancer. reTFAE targets and lung cancer targets share 62 proteins. (B) Protein-Protein-Interaction (PPI) network of 62 proteins. AKT1, TP53, VEGFA, JUN show more connections with other proteins. (C) KEGG Pathway analysis of 62 proteins. P<0.01 was the cutoff for displaying the pathways. (D) The inhibition of PI3K/Akt-suppression and reTFAE on wound healing. (E) The inhibition of PI3K/Akt-suppression and reTFAE on A549 cell invasion. (F) The influence of PI3K/Akt-suppression and reTFAE on the expression of EMT-related proteins. Statistical differences were determined by a two-sided Student’s t-test. Compared with control, N.S., not significant, \*\*\**P* < 0.001; \*\**P* < 0.01; \**P* < 0.05.

Compared with normal cells, inhibition of PI3K/Akt pathway showed poor migration and invasion capabilities (Fig.2D and 2E), and increased expression of E-cadherin and decreased expression of Vimentin, N-cadherin, TWIST1 and p-AKT (Fig.2F), which were related with EMT progress. Therefore, we believed that reTFAE ameliorated the EMT process of A549 cells by inhibiting the PI3K/Akt signaling pathway.

### 3.3. reTFAE inhibited TWIST1 in EMT process of A549 cells

To identify the key target of reTFAE, Cytoscape 3.6.1 was used to construct “Ingredient-Target-Pathway” network. Among proteins involved in the network, TWIST1 attracted our attentions, which was reported to be directly regulated by PI3K-Akt and connected with both baicalein and wogonin[24-27] (Fig.3A). To investigate whether TWIST1 played an important role in reTFAE’s effect on the EMT process, we applied small interfering RNA (siRNA) method to knock down the TWIST1 gene in A549 cell.

**Fig.3.**
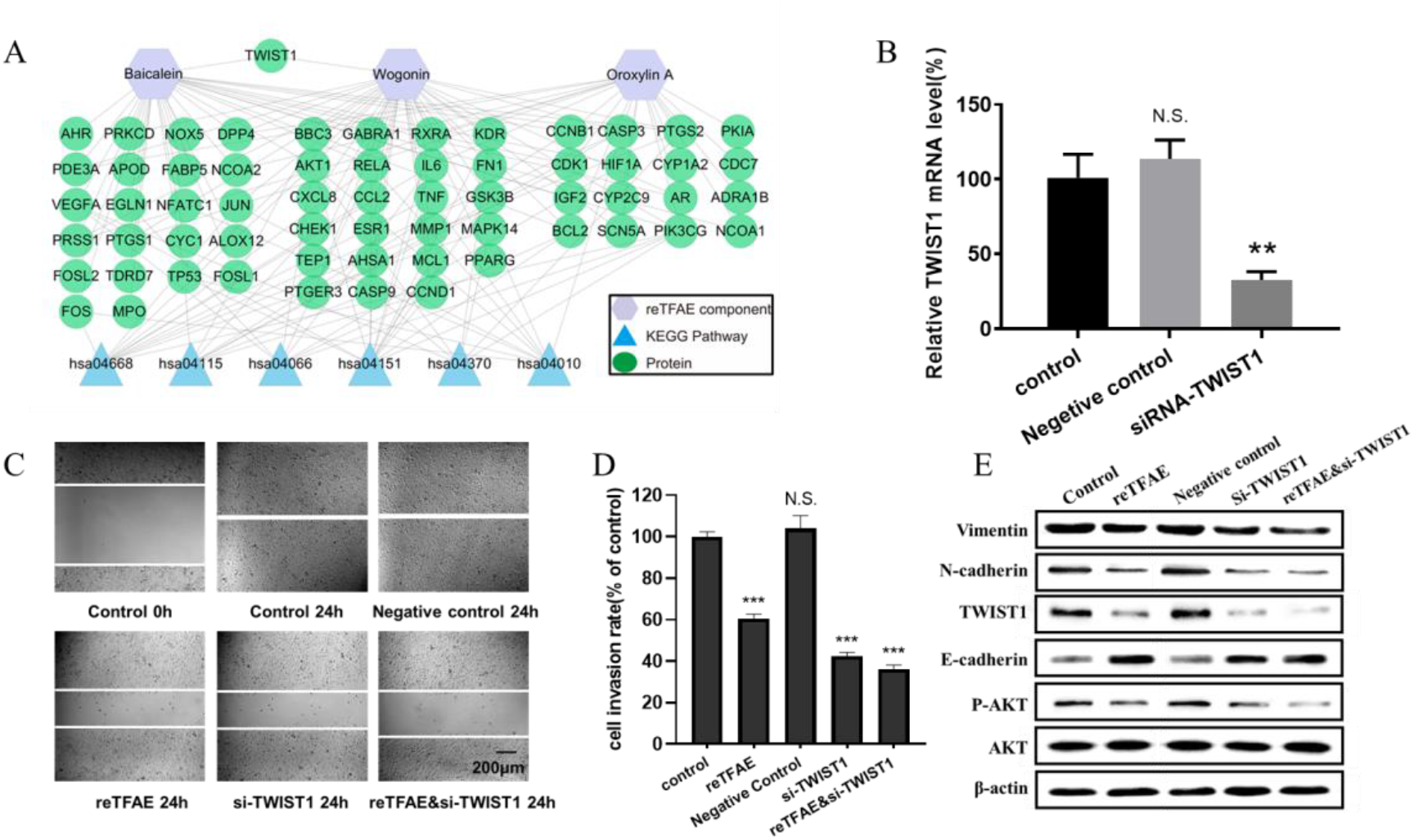
reTFAE ameliorated the EMT process of A549 cells via inhibiting TWIST1. (A)Relationships of reTFAE components-KEGG Pathways-Protein targets. (B)Relative TWIST1 mRNA level after siRNA interference. (C)The inhibitions of siTWIST1 and reTFAE on wound healing. (D)The inhibitions of siTWIST1 and reTFAE on A549 cell invasion rate. (E)The influence of siTWIST1 and reTFAE on the expression of EMT-related proteins. Statistical differences were determined by a two-sided Student’s t-test; Compared with control, N.S., not significant, \*\*\**P* < 0.001; \*\**P* < 0.01; \**P* < 0.05.

Compared with normal cells, TWIST1-knockdowned A549 cells showed significant lower mRNA level (Fig.3B), and reTFAE-treated/ siTWIST1 /reTFAE&siTWIST1-treated cells showed poor migration and invasion capabilities, increased expression of epithelial marker(E-cadherin) and decreased expression of mesenchymal markers (Vimentin and N-cadherin), EMT-related transcription factor (TWIST1) and protein in PI3K/Akt pathway(p-AKT) (Fig.3C and 3D). siTWIST1 showed similar effects to reTFAE, and the synergistic application of reTFAE and siTWIST1 showed more obvious effect (Fig.3E). Therefore, we claimed that reTFAE ameliorated the EMT process of A549 cells via inhibiting TWIST1.

### 3.4. The effect of reTFAE on protein expression of A549 cells

As above, we found that reTFAE inhibited PI3K/Akt pathway and TWIST1, yet how reTFAE led to the decline of EMT is not clear. To explain the further mechanism, we collected A549 cells treated with reTFAE for 24 hours and performed stable-isotope dimethyl labeling proteomics experiment. The proteins of A549 cells treated with DMSO or reTFAE were marked with “light” or “heavy”, respectively, then mixed, desalted, and analyzed by LC-MS/MS(Fig.4A). Proteins with Average Ratio Light/ Heavy(L/H)>1.5 were filtered out, and 158 proteins down-regulated by reTFAE appeared at least twice in repeated experiments (Fig.4B). KEGG pathway analysis was performed on the 158 proteins and reTFAE down-regulated pathways with P<0.01 were as follows: Carbon metabolism, Glycolysis/Gluconeogenesis, Biosynthesis of amino acid and Pentose phosphate pathway (Fig.4C). These four pathways contain 11, 9, 7, and 4 proteins respectively (Table1). In short, reTFAE interferes with the sugar metabolism of A549 cells.

**Fig.4.**
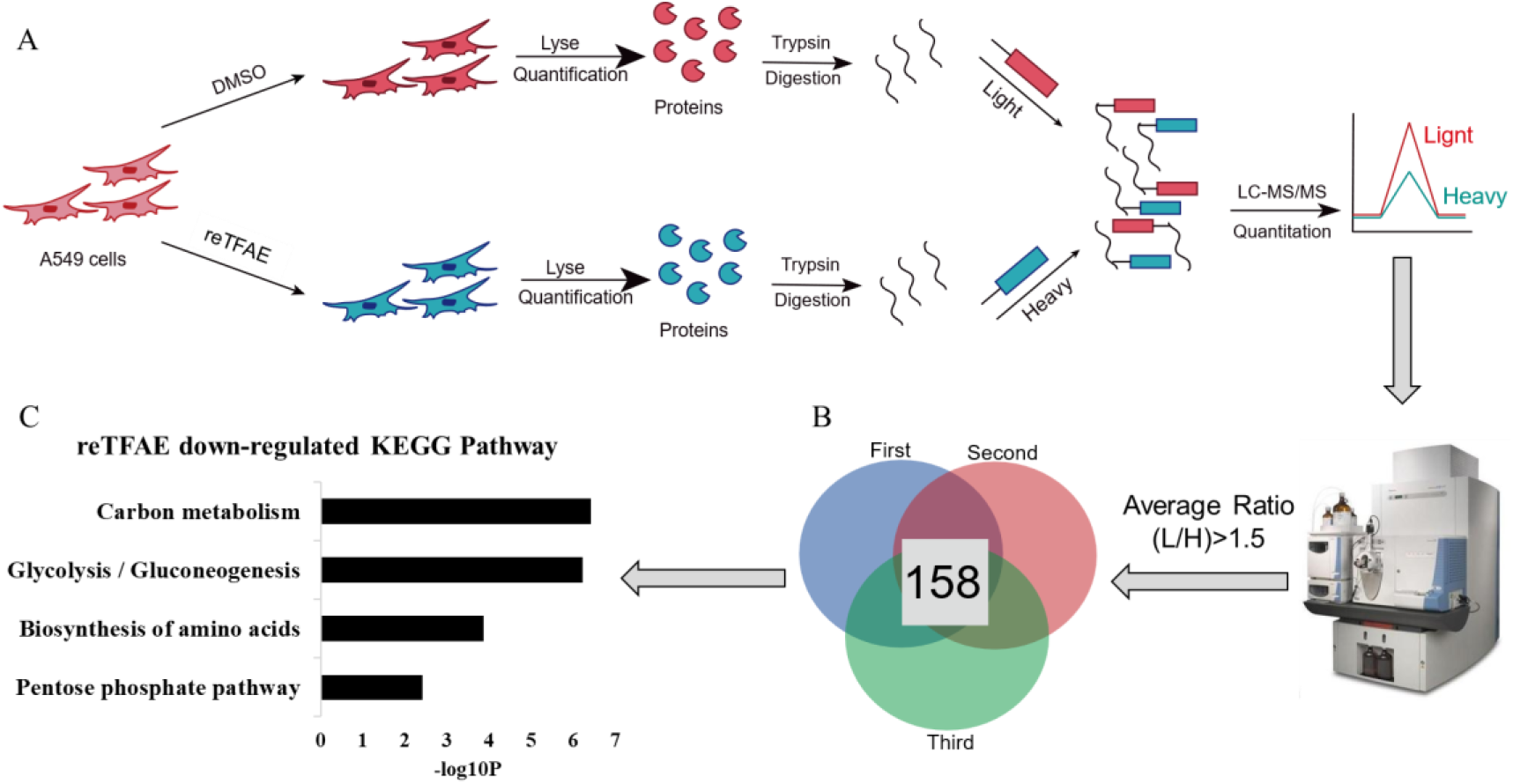
reTFAE interfered with the glucose metabolism of A549 cells. (A)Overall scheme of In-solution dimethyl labeling experiment, the control or reTFAE group were labeled with “light” or “heavy” respectively. (B) Venn diagram showing the 158 proteins with an averaged ratio >1.5 appeared twice of triple experiment. (C)KEGG Pathway analysis of the 158 proteins. P<0.01 was the cutoff for displaying the pathways.

**Table 1.** Pathways interfered by reTFAE, and proteins enriched in each pathway

| Pathways | Enriched Targets |
| --- | --- |
| Carbon metabolism | GPI, PGAM1, PKM, HADHA, TALDO1, GAPDH, PGK1, MDH2, TPI1, ALDOA, G6PD |
| Glycolysis/Gluconeogenesis | GPI, PGAM1, PKM, LDHA, GAPDH, PGK1, TPI1, ALDH3A1, ALDOA |
| Biosynthesis of amino acid | PGAM1, PKM, TALDO1, GAPDH, PGK1, TPI1, ALDOA |
| Pentose phosphate pathway | GPI, TALDO1, ALDOA, G6PD |

## 4. Discussion

As reported, the Total Flavonoid Aglycones Extract of *Scutellaria baicalensis* has an inhibitory effect on NSCLC[23]. In this study, we configured reTFAE and found that it inhibited the EMT process of A549 cells by affecting PI3K/Akt pathway and TWIST1. As “knockdown TWIST1” and “reTFAE incubation” showed similar EMT-inhibition tendency, we believe that reTFAE inhibits PI3K/Akt pathway and TWIST1, thereby inhibiting the EMT process of A549 cells. Actually, some evidence showed that TWIST1-related axis may participate in the EMT process by activating the Wnt/β-catenin signaling pathway, thereby accelerating the process of lung cancer[28]. TWIST1 promotes EMT and metastasis through serine phosphorylation of p38, c-Jun N-terminal kinase (JNK) and Erk1/2[29]. As our research provides direct evidence, TWIST1-targeted flavonoid may provide a new strategy to inhibit EMT progress.

Furthermore, stable-isotope dimethyl labeling proteomics was employed to detail the pharmacodynamic network of lung cancer cells treated with TFAE. In this research, reTFAE down-regulated the carbon metabolism, glycolysis/gluconeogenesis, biosynthesis of amino acid, pentose phosphate pathway, etc. Actually, tumor cells obtain energy mainly through the process of glycolysis, which promotes the EMT process of tumor cells[30]; when nutrients were depleted, cancer cells tend to obtain metabolic materials through the pentose phosphate pathway[31]. Studies had shown that TWIST1 can promote glycolysis process[32-34]. In this study, ALDOA and GPI were enriched in both glycolysis and pentose phosphate pathways, while PKM and LDHA in glycolysis (Table1.). The relationships between the above genes and the EMT process had been reported: PKM expresses pyruvate kinase and catalyzes the transfer of phosphorylation groups from phosphoenolpyruvate to ADP to generate ATP and pyruvate[35]. PKM is expressed in fetal tissues and cancers, and participates in the EMT process of human colon cancer cells [36]. PKM interacts with TGFβ-induced factor homeobox 2 (TGIF2) to inhibit the transcription of E-Cadherin[37]. Glucose-6-phosphate isomerase edited by GPI catalyzes the conversion of glucose-6-phosphate to fructose-6-phosphate. GPI can also be produced by cancer cells to promote the EMT process[38]. GPI promotes the EMT process of breast cancer cells by inhibiting miR-200 and inducing ZEB1/2[39].Silencing GPI promotes the transition of human lung fibroblasts from the mesenchymal to the epithelial state[40].ALDOA expresses fructose-bisphosphate aldolase A, which is involved in glycolysis and gluconeogenesis, and is also overexpressed in cancer[41]. ALDOA is highly expressed in lung squamous cell carcinoma (LSCC) and depletion of ALDOA in lung squamous carcinoma cells reduces cell motility capabilities [42]. Overexpression of ALDOA in colon cancer cells leads to the EMT progress[43].In pancreatic cancer and bladder cancer cells, ALDOA-silencing increases E-Cadherin and decrease N-Cadherin expression[44, 45]. Lactate dehydrogenase-A (LDHA) enzyme converts pyruvate into lactic acid. LDHA promotes the EMT process, but the specific mechanism is not clear[46, 47]. our hypothesis is that reTFAE had a relationship with glycolysis or PI3K/Akt-related proteins to exert EMT-inhibiting effect, yet it needs further verification.

Moreover, we also explored the effects of the three main components of reTFAE-baicalein, wogonin, and oroxylin-A on the EMT process, and found that baicalein had the most significant inhibitory effect on the EMT process of A549 cells, closing to the effect of reTFAE. Next, we will explore the inhibitory mechanism of baicalein on NSCLC.

## 5. Conclusion

In this work, we first determined that reTFAE composed of three compounds in a definite proportion had the activity of inhibiting the invasion and metastasis of lung cancer A549 cells. Then combing with network pharmacology and molecular biology technology, PI3K-AKT-TWIST1 axis was inhibited by reTFAE to ameliorate EMT progress. Furthermore, chemoproteomics was used to elucidate the changes of downstream protein networks after TFEA inhibited PI3K-AKT-TWIST1 axis, and it was found that glycolysis pathway may play an important role. In summary, reTFAE can inhibit the PI3K/Akt pathway with the core of TWIST1, thereby inhibiting the glycolytic pathway to suppress EMT in A549 cells (Fig.5). TWIST1-targeted flavonoid provided a new strategy to inhibit EMT progress for the treatment of NSCLC.

**Fig.5.**
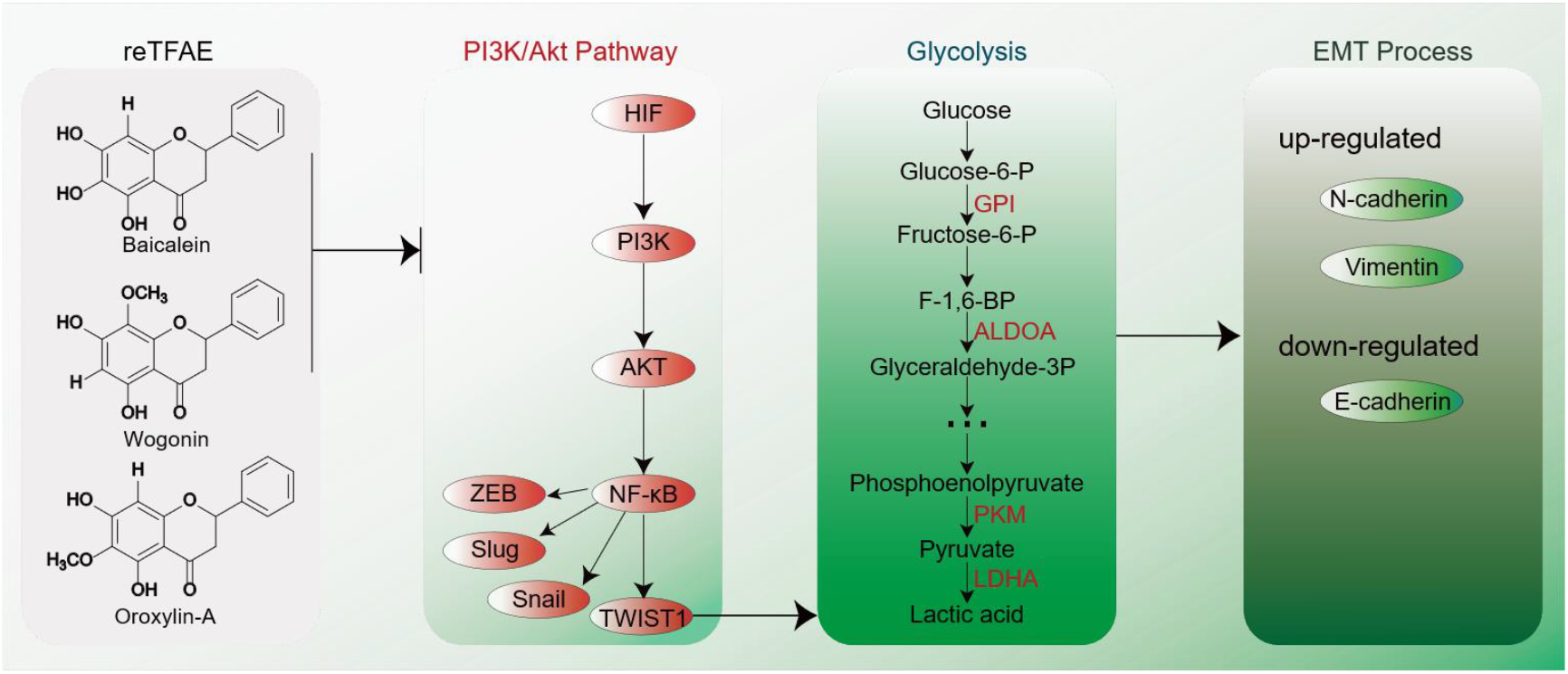
EMT process induced by reTFAE. Baicalein and wogonin inhibit the PI3K/Akt pathway, thereby inhibiting TWIST1, which inhibits the glycolysis pathway and inhibits the EMT process of A549 cells.

## 6. Acknowledgement

This work is funded by the Science and Technology Project of Hebei Education Department (QN2018060), National Key Research and Development Program of China (2018YFC1706300 and 2018YFC17063005), the Science and Technology Planning Project of Gansu Province (20JR10RA586), the Fundamental Research Funds for the Central Universities(lzujbky-2021-kb40), the Project for Longyuan Youth Innovation and Entrepreneurship Talent.

## References

[1] 2020 World Health Statistics. 2020 2020; Available from: https://www.who.int/data/collections.

[2] Mph, R.L.S., et al., Cancer Statistics, 2017. Ca-Cancer J. Clin., 2017. 67: p. 7–30.

[3] Lopez-Lazaro, M., The Warburg Effect: Why and How Do Cancer Cells Activate Glycolysis in the Presence of Oxygen? Anti-Cancer Agents Med. Chem., 2008. 8: p. 305–312.

[4] Gatenby, R.A. and R.J. Gillies, Why do cancers have high aerobic glycolysis? Nat. Rev. Cancer, 2004. 4: p. 891–899.

[5] Heiden, M.G.V., L.C. Cantley, and C.B. Thompson, Understanding the Warburg Effect: The Metabolic Requirements of Cell Proliferation. Science, 2009. 324: p. 1029–1033.

[6] Muz, B., et al., The role of hypoxia in cancer progression, angiogenesis, metastasis, and resistance to therapy. Hypoxia, 2015. 2015: p. 83–92.

[7] Eales, K.L., K.E.R. Hollinshead, and D.A. Tennant, Hypoxia and metabolic adaptation of cancer cells. Oncogenesis, 2016. 5: p. e190.

[8] Keith, R.L., Lung carcinoma. 2017(https://www.msdmanuals.com/professional/pulmonary-disorders/tumors-of-the-lungs/lung-carcinoma.%20Accessed%20Nov%208,%202017.).

[9] Tulchinsky, E., et al., EMT: A mechanism for escape from EGFR-targeted therapy in lung cancer. BBA-Rev. Cancer, 2018. 1871: p. 29–39.

[10] T, S.A. and S. Benjamin, Targeting anaplastic lymphoma kinase in lung cancer. Clin. Cancer Res., 2011. 17: p. 2081–2086.

[11] Otsuki, Y., H. Saya, and Y. Arima, Prospects for new lung cancer treatments that target EMT signaling. Dev. Dyn., 2018. 247: p. 462–472.

[12] Tieju, L., et al., The EMT transcription factor, Twist1, as a novel therapeutic target for pulmonary sarcomatoid carcinomas. Int. J. Mol. Sci., 2020. 56: p. 750–760.

[13] Jente, v.S., et al., Epithelial-mesenchymal-transition-inducing transcription factors: new targets for tackling chemoresistance in cancer? Oncogene, 2018. 37: p. 6195–6211.

[14] L, S.G., HIF-1: upstream and downstream of cancer metabolism. Curr. Opin. Genet. Dev., 2010. 20: p. 51–56.

[15] Tennant, D.A., R.V. Durán, and E. Gottlieb, Targeting metabolic transformation for cancer therapy. Nat. Rev. Cancer, 2010. 10: p. 267–277.

[16] Gongda, X., et al., Akt/PKB-mediated phosphorylation of Twist1 promotes tumor metastasis via mediating cross-talk between PI3K/Akt and TGF-β signaling axes. Cancer Discovery, 2012. 2: p. 248–259.

[17] Ding, X., F. Li, and L. Zhang, Knockdown of Delta-like 3 restricts lipopolysaccharide-induced inflammation, migration and invasion of A2058 melanoma cells via blocking Twist1-mediated epithelial-mesenchymal transition. Life Sci., 2019. 226: p. 149–155.

[18] Hong, R., et al., TWIST1 and BMI1 in Cancer Metastasis and Chemoresistance. J. Cancer, 2016. 7: p. 1074–1080.

[19] Qing-Qing, Z., et al., The role of TWIST1 in epithelial-mesenchymal transition and cancers. Tumor Biol., 2016. 37: p. 185–197.

[20] Yochum, Z.A., et al., Targeting the EMT transcription factor TWIST1 overcomes resistance to EGFR inhibitors in EGFR-mutant non-small-cell lung cancer. Oncogene, 2019. 38: p. 656–670.

[21] P, C., et al., Plasma concentrations of VCAM-1 and ICAM-1 are elevated in patients with Type 1 diabetes mellitus with microalbuminuria and overt nephropathy. Diabetic Med., 2000. 17: p. 644–649.

[22] Dorota, W., D. Andrzej, and M. Adam, Antiradical and antioxidant activity of flavones from Scutellariae baicalensis radix. Nat. Prod. Res., 2015. 29: p. 1567–1570.

[23] Yang, W., et al., Total flavonoid aglycones extract in Radix scutellariae inhibits lung carcinoma and lung metastasis by affecting cell cycle and DNA synthesis. J. Ethnopharmacol., 2016. 194: p. 269–279.

[24] Yuan, R., J. Chang, and J. He, Effects of Twist1 on drug resistance of chronic myeloid leukemia cells through the PI3K/AKT signaling pathway. Cellular and molecular biology (Noisy-le-Grand, France), 2020. 66(6): p. 81.

[25] Xue, G., et al., Akt/PKB-Mediated Phosphorylation of Twist1 Promotes Tumor Metastasis via Mediating Cross-Talk between PI3K/Akt and TGF-β Signaling Axes. Cancer Discovery, 2012. 2(3): p. 248–259.

[26] Zeng, et al., Baicalein suppresses the proliferation and invasiveness of colorectal cancer cells by inhibiting Snailinduced epithelialmesenchymal transition. Molecular Medicine Reports, 2020.

[27] Yao, Y., et al., Wogonoside inhibits invasion and migration through suppressing TRAF2/4 expression in breast cancer. Journal of Experimental & Clinical Cancer Research, 2017. 36(1): p. 103.

[28] Pan, J., et al., lncRNA JPX/miR-33a-5p/Twist1 axis regulates tumorigenesis and metastasis of lung cancer by activating Wnt/β-catenin signaling. Mol. Cancer, 2020. 19.

[29] Jun, H., et al., Phosphorylation of serine 68 of Twist1 by MAPKs stabilizes Twist1 protein and promotes breast cancer cell invasiveness. Cancer Res., 2011. 71: p. 3980–3990.

[30] Liberti, M.V. and J.W. Locasale, Correction to: ‘The Warburg Effect: How Does it Benefit Cancer Cells?’. Trends Biochem. Sci., 2016. 41: p. 211–218.

[31] Ilias, G.-S., et al., EMT Factors and Metabolic Pathways in Cancer. Front Oncol, 2020. 10.

[32] Sara, L., et al., Endothelial-to-mesenchymal transition compromises vascular integrity to induce Myc-mediated metabolic reprogramming in kidney fibrosis. Sci. Signaling, 2020. 13: p. eaaz2597.

[33] Li, Y., et al., Twist promotes reprogramming of glucose metabolism in breast cancer cells through PI3K/AKT and p53 signaling pathways. Oncotarget, 2015. 6: p. 25755–25769.

[34] Wang, X.-X., et al., TWIST1 transcriptionally regulates glycolytic genes to promote the Warburg metabolism in pancreatic cancer. Exp. Cell Res., 2020. 386: p. 111713.

[35] Dombrauckas, J.D., B.D. Santarsiero, and A.D. Mesecar, Structural basis for tumor pyruvate kinase M2 allosteric regulation and catalysis. Acta Crystallogr., Sect. A: Found. Crystallogr., 2005. 61: p. 9417–9429.

[36] Wong, N., et al., PKM2 contributes to cancer metabolism. Cancer Lett., 2015. 356: p. 184–191.

[37] Hamabe, A., et al., Role of pyruvate kinase M2 in transcriptional regulation leading to epithelial– mesenchymal transition. Proc. Natl. Acad. Sci., 2014. 111: p. 15526–15531.

[38] Y, N., et al., Expression and secretion of neuroleukin/phosphohexose isomerase/maturation factor as autocrine motility factor by tumor cells. Cancer Res., 1998. 58: p. 2667–2674.

[39] Tatsuyoshi, F., H. Victor, and R. Avraham, Phosphoglucose isomerase/autocrine motility factor mediates epithelial and mesenchymal phenotype conversions in breast cancer. Cancer Res., 2009. 69: p. 5349–5356.

[40] Tatsuyoshi, F., et al., Down-regulation of phosphoglucose isomerase/autocrine motility factor results in mesenchymal-to-epithelial transition of human lung fibrosarcoma cells. Cancer Res., 2007. 67: p. 4236–4243.

[41] Yu-Chan, C., et al., Roles of Aldolase Family Genes in Human Cancers and Diseases. Trends Endocrinol. Metab., 2018. 29: p. 549–559.

[42] Du, S., et al., Fructose-Bisphosphate Aldolase A Is a Potential Metastasis-Associated Marker of Lung Squamous Cell Carcinoma and Promotes Lung Cell Tumorigenesis and Migration. PLoS One, 2014. 9: p. e85804.

[43] Feng, Y., et al., Aldolase A overexpression is associated with poor prognosis and promotes tumor progression by the epithelial-mesenchymal transition in colon cancer. Biochem. Biophys. Res. Commun., 2018. 497: p. 639–645.

[44] Ji, S., et al., ALDOA functions as an oncogene in the highly metastatic pancreatic cancer. Cancer Lett., 2016. 374: p. 127–135.

[45] Jianwei, L., et al., ALDOLASE A regulates invasion of bladder cancer cells via E-cadherin-EGFR signaling. J. Cell. Biochem., 2019: p. 13694–13705.

[46] Tirpe, A.A., et al., Hypoxia: Overview on Hypoxia-Mediated Mechanisms with a Focus on the Role of HIF Genes. Int. J. Mol. Sci., 2019. 20: p. 6140.

[47] Hou, X.-m., et al., LDH-A promotes malignant behavior via activation of epithelial-to-mesenchymal transition in lung adenocarcinoma. Biosci. Rep., 2019. 39: p. BSR20181476.

