## Supplementary figures and images for "The compound of baicalein, wogonin, and oroxylin-A inhibits EMT in A549 cell line via PI3K/AKT-TWIST1- glycolysis pathway"

### Supplementary Information

#### Supporting Information

##### Western blot pictures:

##### 1.TWIST1:

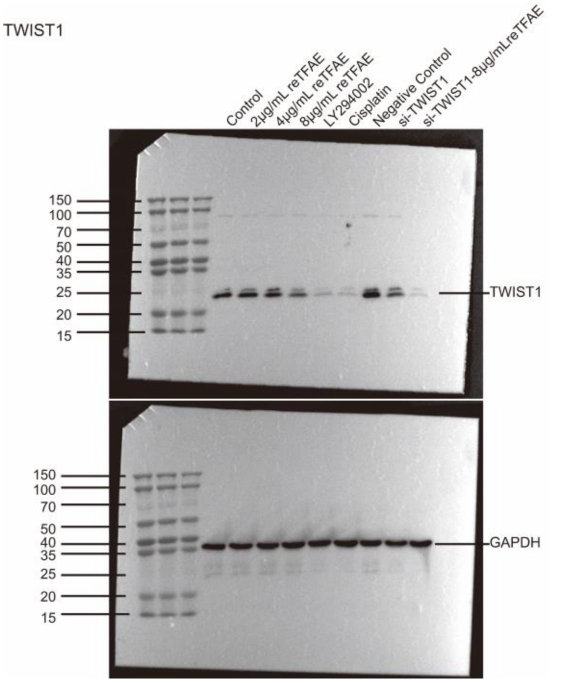


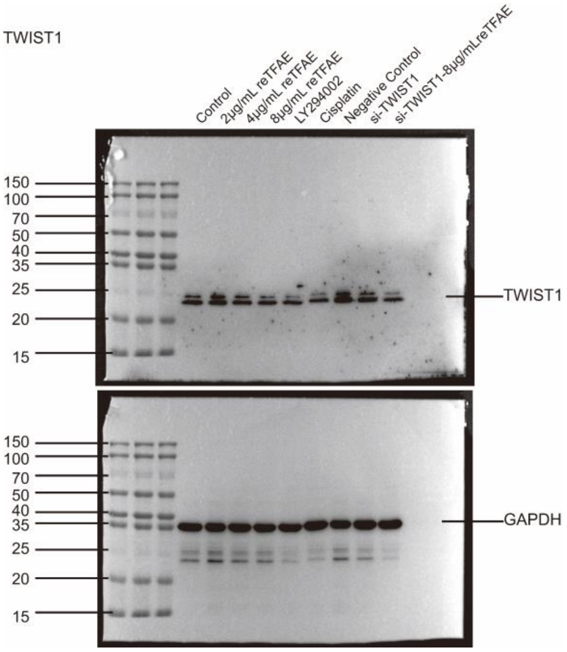

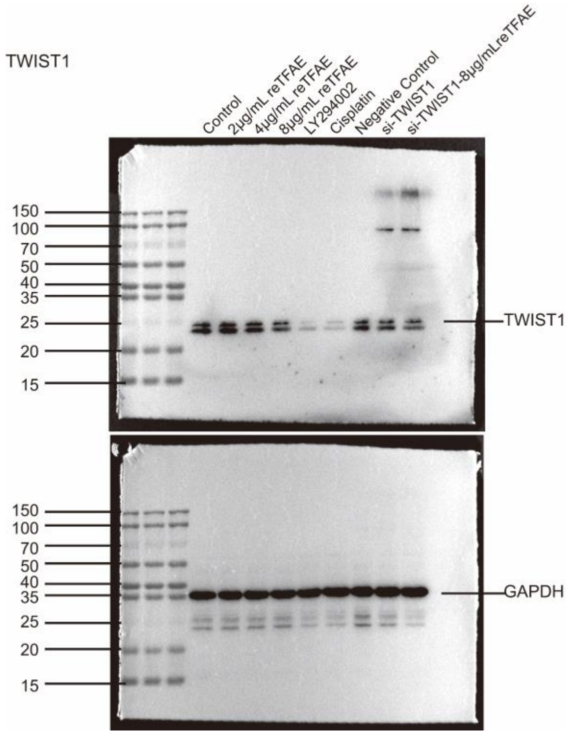

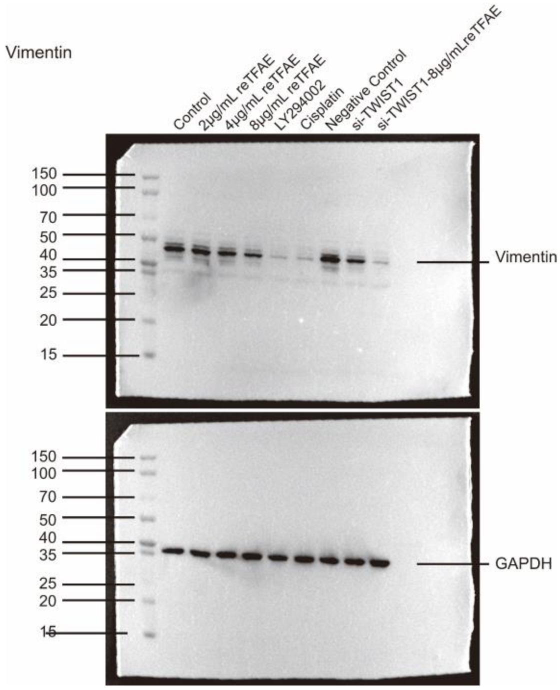

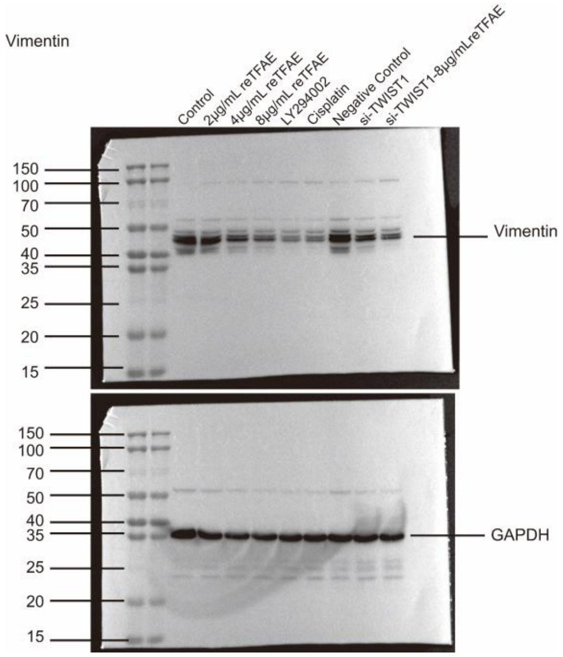

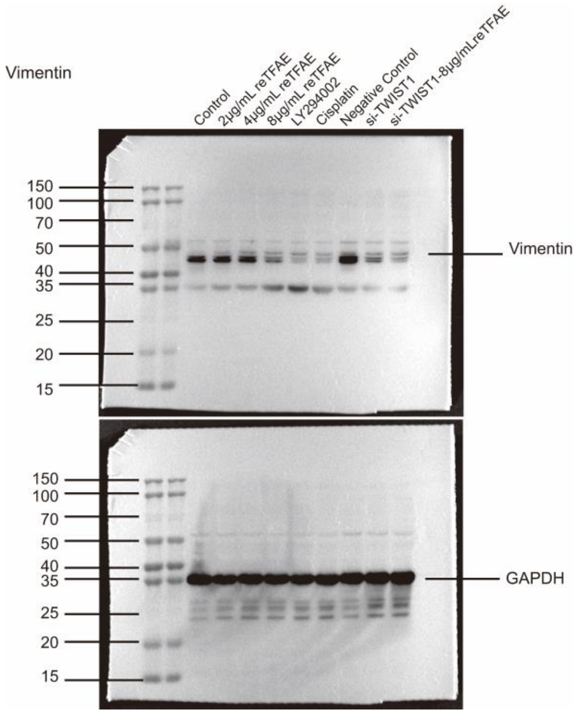

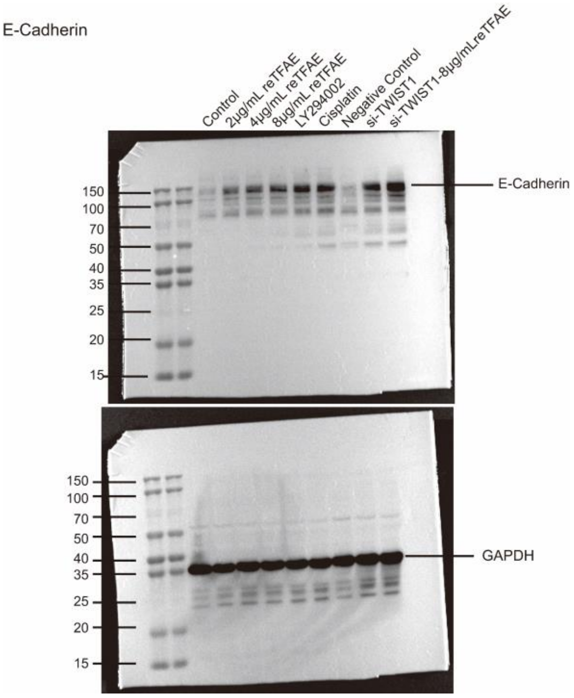

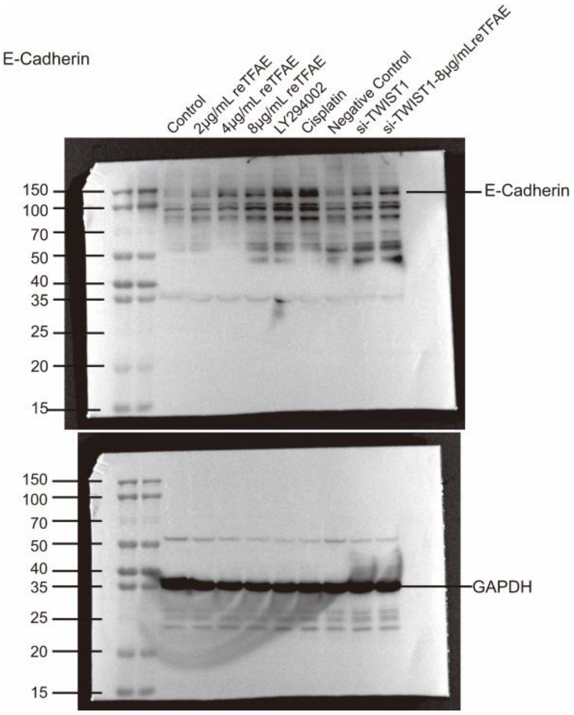

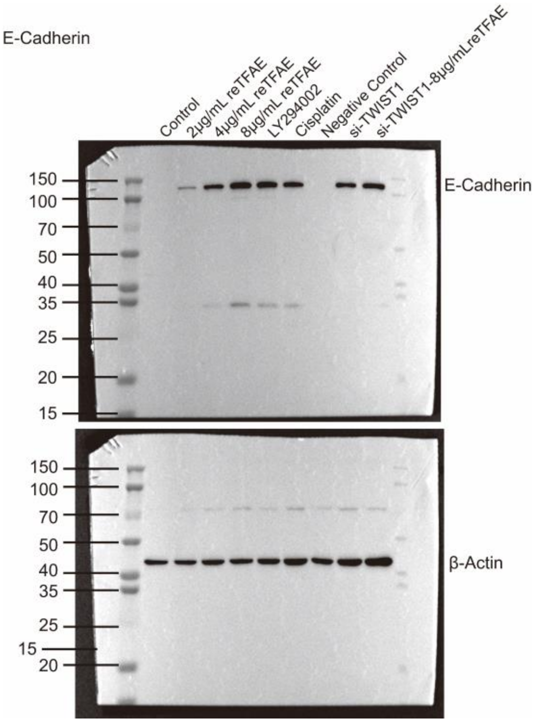

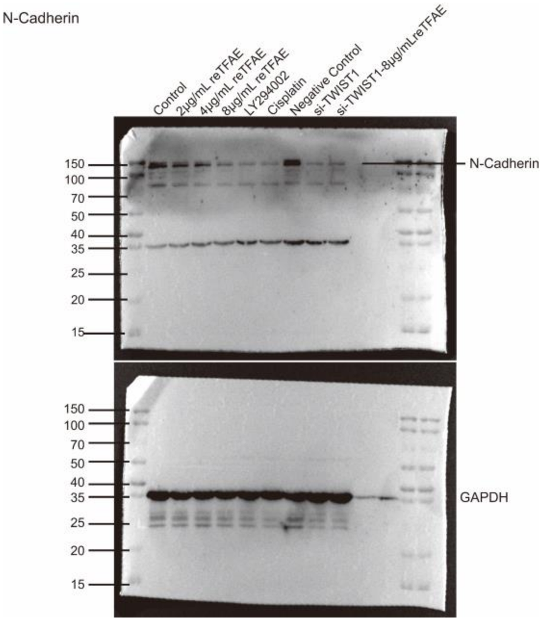

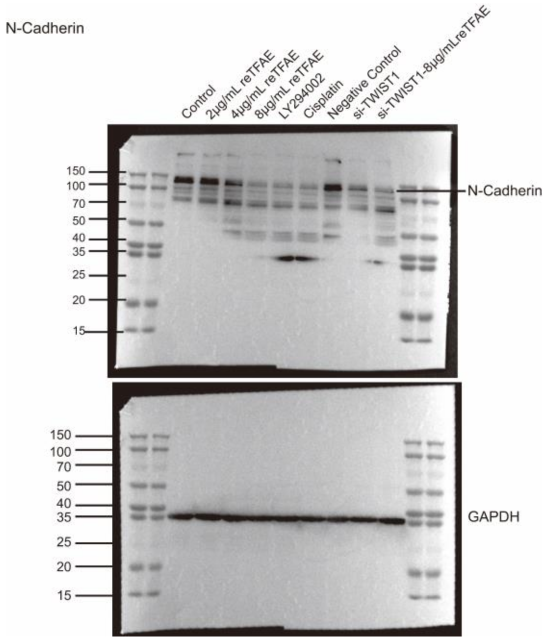

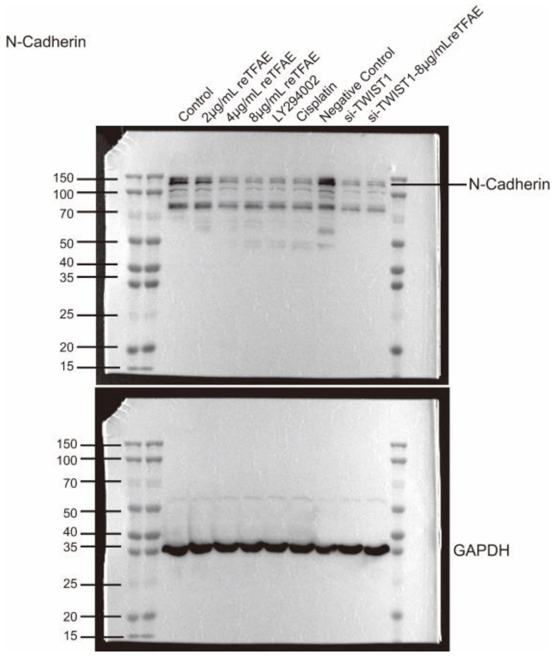

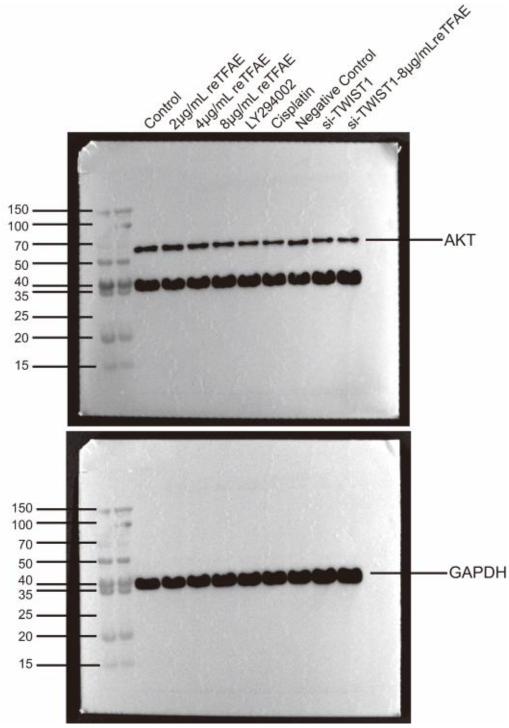

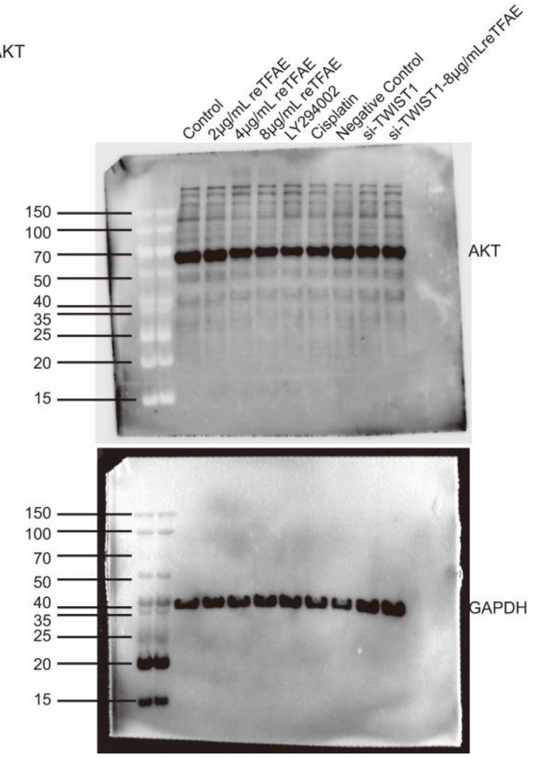

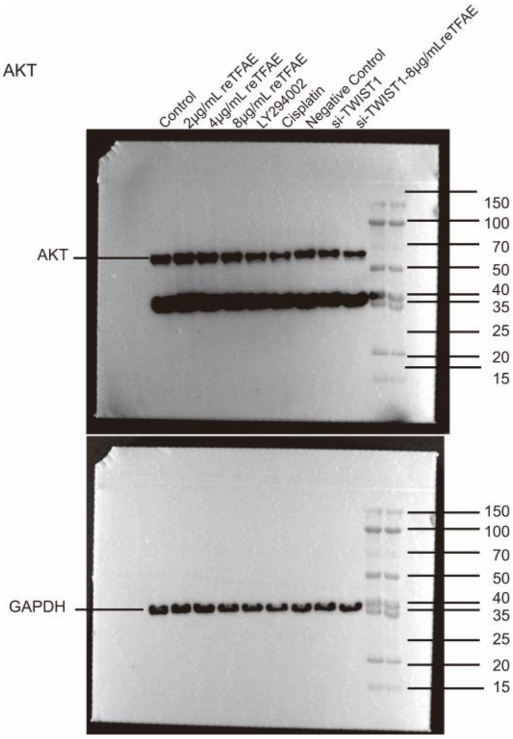

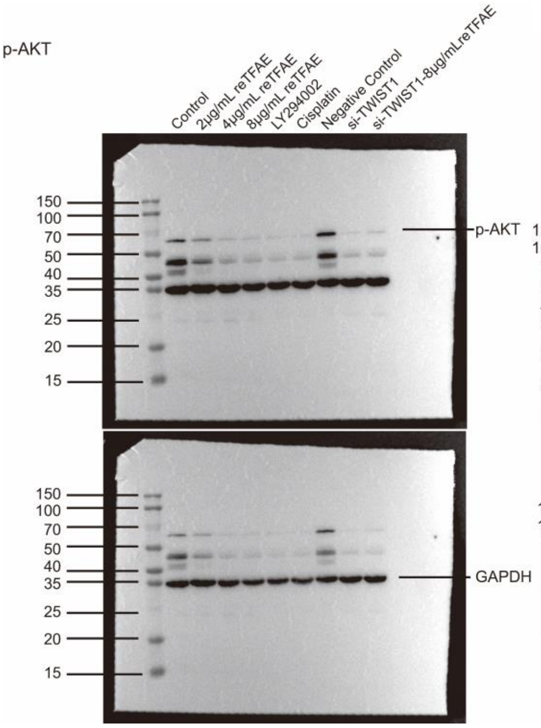

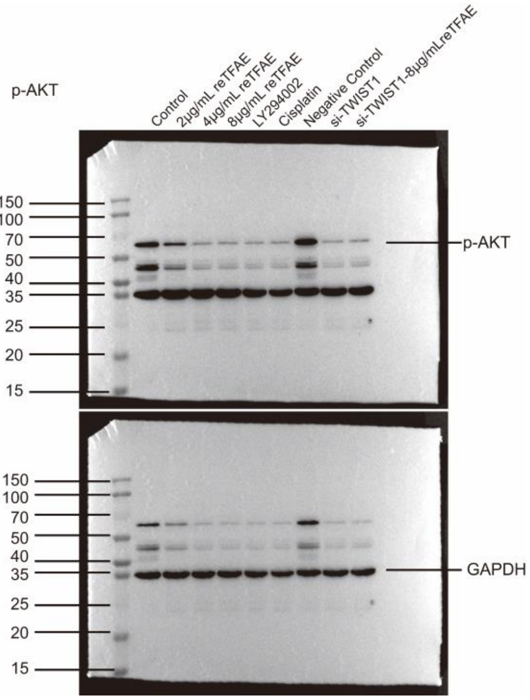

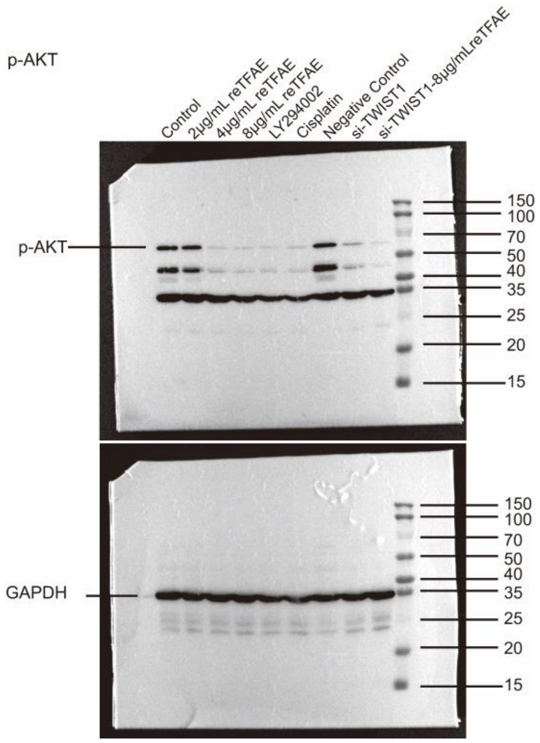
